# Trem-2 promotes emergence of restorative macrophages and endothelial cells during recovery from hepatic tissue damage

**DOI:** 10.1101/823773

**Authors:** Inês Coelho, Nádia Duarte, André Barros, Maria Paula Macedo, Carlos Penha-Gonçalves

**Author notes:** Corresponding author: Carlos Penha-Gonçalves, Address: Instituto Gulbenkian de Ciência (IGC), Rua da Quinta Grande n°6, 2780-156 Oeiras, Portugal.

## Abstract

Macrophages are pivotal in mounting liver inflammatory and tissue repair reactions upon hepatic injury showing remarkable functional plasticity. Nevertheless, the molecular mechanisms determining macrophage transition from inflammatory to restorative phenotypes in the damaged liver remain unclear. Using mouse models of acute (APAP) or chronic (CCl4) drug-induced hepatotoxic injury we show that the immune receptor Trem-2 controls phenotypic shifts in liver macrophages and impacts endothelial cell differentiation during tissue recovery.

Trem-2 gene ablation led to delayed re-population of Kupffer cells correlating with deterred resolution of hepatic damage following acute and chronic injury. We found that during tissue recovery macrophages in transition to the Kupffer cell compartment expressed high levels of Trem-2. Acquisition of the transition phenotype was associated with an unique transcriptomic profile denoting strong responsiveness to oxidative stress and downmodulation of the pro-inflammatory phenotype, which was not observed in absence of Trem-2.

During tissue recovery lack of Trem-2 favored accumulation of a liver-damage associated endothelial cell population (LDECs) engaged in a transcriptional program compatible with endothelial de-differentiation. Accordingly, LDECs precursor potential is supported by the downregulation of surface endothelial cell markers and striking *in vitro* morphological changes towards typical endothelial cells.

In conclusion, we found that the dynamics of liver macrophages in response to liver injury is critically controlled by Trem-2 and is interlinked with the de-differentiation of endothelial cells and heightened liver pathology. We propose that Trem-2 promotes the transition from the pro-inflammatory to the tissue repair phase by driving the acquisition of restorative properties of phagocytic macrophages.

## Introduction

Hepatotoxic insults elicit a multilayered response involving clearance of damaged tissue, scar formation and tissue regeneration. Macrophages play decisive roles in inflammatory and tissue repair responses during acute and chronic liver injury^1–3^ and in liver damage associated to metabolic disorders such as NAFLD^4,5^, type 2 diabetes and obesity^6^.

In the damaged hepatic tissue macrophages with different surface phenotypes and activation status show sharp population dynamics^1,3^ suggesting that distinct macrophage populations perform specific activities and are determinants of the course of response to tissue damage. The macrophage involvement in the response to severe damage is often initiated by influx of hematopoietic-derived monocytes that home the liver as Ly6c^+^ cells and dominate the liver macrophage populations at this stage^2,3,7^. These cells present a high-inflammatory phenotype including the expression of TNF-α, IL-1β and TGF-β that amplify tissue pathology and under chronic tissue injury promote transdifferentiation of stellate cells into collagen-producing myofibroblasts, the hallmark of liver fibrosis^1^.

On the other hand, tissue repair and fibrosis resolution are associated with the emergence of macrophages that phenotypically resemble liver resident macrophage cells^1,3^. These pro-resolution macrophages show phagocytic ability and express high levels of metalloproteinases and anti-inflammatory mediators (e.g. MMP12 and Arg1) that trigger myofibroblasts apoptosis and actively participate in extracellular matrix degradation^1^. Nevertheless, it is unclear what controls the dynamics of different liver macrophage populations during response to damage. Macrophages show remarkable phenotypic and functional plasticity and are equipped to undergo functional transitions, depending on contextual cues^8,9^. Interestingly, it has been shown that pro-inflammatory macrophages can acquire anti-inflammatory and pro-repair phenotypes^1,3,10^ but the triggers of this phenotype switch in liver macrophages remains largely unknown^1,9^.

Triggering receptor expressed on myeloid cells-2 (Trem-2) is a transmembrane immune receptor typically expressed in the monocyte/macrophage lineage^11^. Upon ligand binding Trem-2 signals through the adaptor DAP12 impacting on activation of macrophage effector functions^12^. Trem-2 has been intensively studied in the context of neurodegenerative diseases, that revealed its role in engulfment of αβ-amyloid plaques during Alzheimer’s disease^13^ and phagocytosis of apoptotic neurons^14^. In addition, Trem-2 signaling was shown to limit tissue destruction and to facilitate repair, clearing cellular debris in a model of Experimental Autoimmune Encephalomyelitis (EAE)^15^. Trem-2 ligands leading to macrophage activation *in situ* have not been identified but different studies reported the binding to phospholipids such as phosphatidylserine^14,16^ and a range of acidic and zwitterionic lipids^13^, which may accumulate upon cell damage.

In addition, Trem-2 has been shown to modulate microglia survival through Wnt/β-catenin signaling^17,18^ and to promote inhibitory signals that restrain pro-inflammatory macrophage activation^19,20^. Recent studies show that Trem-2 is required for activation of a specific transcriptional gene program which controls phagocytosis and lipid metabolism of microglial cells in Alzheimer’s disease^21^ and of lipid associated macrophages (LAM) in metabolic disorders^22^.

The role of Trem-2 in liver macrophages is less explored. Previous studies from our laboratory showed that Trem-2 is expressed on Kupffer cells (KCs) determining their activation profile upon contact with malaria parasite^23^. Recently published work^24^ revealed that Trem-2 is involved in liver damage and proposes that Trem-2 expression in non-parenchymal acts as a brake of the inflammatory response during hepatotoxic injury. Although this established a role for Trem-2 in liver inflammation, its specific effects on macrophages, which are critical players in these processes, remains unsettled.

Here we uncovered that upon experimental induction of hepatic injury Trem-2 controls the dynamics of liver macrophage populations favoring the emergence of restorative macrophages and consequently promoting tissue damage resolution and regeneration of the hepatic tissue, including the endothelial cell lineage.

## Materials and Methods

### Mice and experimental models

All procedures involving laboratory mice were in accordance with national (Portaria 1005/92) and European regulations (European Directive 86/609/CEE) on animal experimentation and were approved by the Instituto Gulbenkian de Ciência Ethics Committee and the Direcção-Geral de Veterinária (the Official National Entity for regulation of laboratory animals usage). Trem2-deficient mice^19^ were kindly provided by Marco Colonna, Washington University School of Medicine, St. Louis, MO. C57BL/6 mice, Trem-2 KO and B6.Actin-GFP mice were bred and housed under a 12-hr light/dark cycle in specific pathogen free housing facilities at the Instituto Gulbenkian de Ciência.

To induce acute liver injury, C57BL/6 and Trem-2 KO male mice with 10 weeks of age were fasted for 15 hours prior to intra-peritoneal injection with 300mg/Kg of acetaminophen (N-acetyl-p-aminophenol (APAP))^3^ (Sigma, St. Louis, MO, USA) in PBS or PBS only. Liver and blood were collected at day 1 (D1) and 3 (D3) after injection. In the model of chronic liver fibrosis and fibrosis regression, C57BL/6 and Trem-2 KO males with 7-8 weeks of age received PBS or 20%v/v carbon tetrachloride (CCl4, Sigma, St. Louis, MO, USA) in olive oil, administered at 0,4mL/Kg, twice a week during 4 weeks by intra-peritoneal injections^1^. Liver and blood were collected at day 1 (fibrosis) or day 3 post-injection (fibrosis regression).

For the *in vivo* phagocytosis experiments, mice were given a retro-orbital injection of 50×10^6^ beads/200uL fluorescent beads (Fluoresbrite YG microspheres 2μm, Polysciences) 3 days after APAP treatment and 1 hour prior liver collection.

### Non-parenchymal cells isolation, flow cytometry and cell sorting

Non-parenchymal cells (NPCs) were isolated from liver lobes by perfusion with Collagenase H (Sigma, St. Louis, MO, USA) followed by density centrifugation as previously described^25,26^. Non-parenchymal cells (NPCs) were immuno-labeled with fluorochrome-conjugated antibodies (eBiosciences and BioLegend) followed by flow cytometry analysis (LSR Fortessa X20™, BD) or cell sorting (FACSAria, BD). Antibodies listed in supplementary methods.

### Histology

Histological analyses were performed in the Histopathology Unit of Instituto Gulbenkian de Ciência. Livers were fixed in 10% formalin and embedded in paraffin. Non-consecutive 3µm sections were stained with hematoxylin-eosin or Mason’s Trichrome and examined under light microscope (Leica DM LB2, Leica Microsystems, Wetzlar, Germany). Necrosis and fibrosis were blindly assessed by a trained pathologist. Necrosis scoring: 0, no necrosis; 1, single cell centrilobular necrosis to centrilobular necrosis without central to central bridging necrosis; 2, centrilobular necrosis with central to central bridging; 3, centrilobular necrosis with central to central bridging and focal coalescent foci of necrosis; 4, centrilobular necrosis with central to central bridging with large coalescent foci of necrosis. Fibrosis was assessed using Ishak system adaptation: 0, no fibrosis; 1, fibrous expansion in some centrilobular areas with or without fibrous septa; 2, fibrous expansion of most centrilobular areas with or without fibrous septa; 3, fibrous expansion of most centrilobular areas with occasional central to central bridging; 4, fibrous expansion in most centrilobular areas with marked central to central bridging; 5, marked bridging (C-C) with occasional nodules; 6: cirrhosis. Images were acquired on a Leica DM LB2 and a commercial Leica High Content Screening microscope.

For immunofluorescence, PFA fixed cells were stained overnight at 4°C with rat anti-mouse F4/80 diluted 1:50 and rabbit anti-mouse caspase-3, diluted 1:500. On the following day sections were washed and incubated with respective secondary antibodies for 1h at room temperature. Images were acquired using a 5×5 tile-scan protocol on a Nikon Ti microscope using a 20× 0.75 NA objective, coupled with an Andor Zyla 4.2 sCMOS camera (Andor, Oxford Instruments) and controlled through Nikon NIS Elements (Nikon). DAPI, F4/80 and caspase-3 were acquired using a DAPI, Cy5 and TRITC filtersets, respectively. Detailed methods for analysis can be found in supplementary methods.

### AST/ALT

Serum levels of Aspartate aminotransferase (AST) and Alanine aminotransferase (ALT) were determined by a colorimetric enzymatic assay using the GOT-GPT kit (Spinreact S.A., Spain) according to manufactures’ instructions.

### RNA isolation and gene expression analysis

NPCs were collected in lysis buffer (RNeasy MiniKit-Qiagen) and total RNA was obtained using RNeasy MiniKit (Qiagen) and converted to cDNA (Transcriptor First Strand cDNA Synthesis Kit, Roche). Cells-to-ct kit (Applied Biosystems) was used to amplify cDNA from sorted cells. Taqman gene expression assays (TNFa Mm00443258_m1, trem2 Mm00451744_m1, Applied Biosystems) and endogenous control GAPDH were used in multiplex Real-Time PCR reactions (ABI QuantStudio-384, ThermoFischer). Results represent relative quantification calculated using the 2-ΔΔCT method and normalized to GAPDH.

### RNA Sequencing

#### Cell sorting and RNA sequencing analysis

Macrophage populations and CD45^neg^ SSC^hi^ population were sorted directly into Qiagen RLT lysis buffer. Each sample represents a pooling of 4 mice. Biological replicates were used for each population except for KCs control. Sequencing was performed at the Genomics Unit, Instituto Gulbenkian de Ciência (IGC, Portugal) following a previously established protocol^27^. Briefly, RNA was separated from gDNA using a modified oligo-dT bead-based strategy and DNA libraries were prepared using Pico Nextera protocol. Sequencing was performed using NextSeq500-High Output Kit v2 (75 cycles), single-ended, 20 million reads per sample. Detailed methods for analysis can be found in supplementary methods.

### Statistical Analysis

GraphPad Prism version 6 software was used to perform statistical analysis.Additional materials and methods are available as supplementary methods.

## Results

### Trem-2 ablation deters tissue repair upon acute liver injury

To study the role of Trem-2 in responses to acute liver injury we used a well-established experimental model^3^. Mice received a single dose of acetaminophen (APAP) and were analyzed after 1 day (D1) or 3 days (D3), corresponding to the times of hepatic damage and tissue repair responses, respectively **(Fig. 1A)**. During acute injury (D1) wild-type and Trem-2 KO mice showed regions of massive necrosis in the liver **(Fig. 1B and C)** but at D3 wild-type had almost completely cleared necrosis while Trem-2 KO mice retained marked liver pathology with wide coalescent necrotic areas **(Fig. 1B and C)**. Serum levels of hepatic enzymes AST and ALT indicated that the extent of APAP hepatotoxicity was not affected by Trem-2 expression **(Fig. 1D and E)**. These data show that although wild-type and Trem-2 KO mice were similarly affected by acute liver injury, resolution of liver damage in Trem-2 KO mice was impaired.

**Figure 1.**
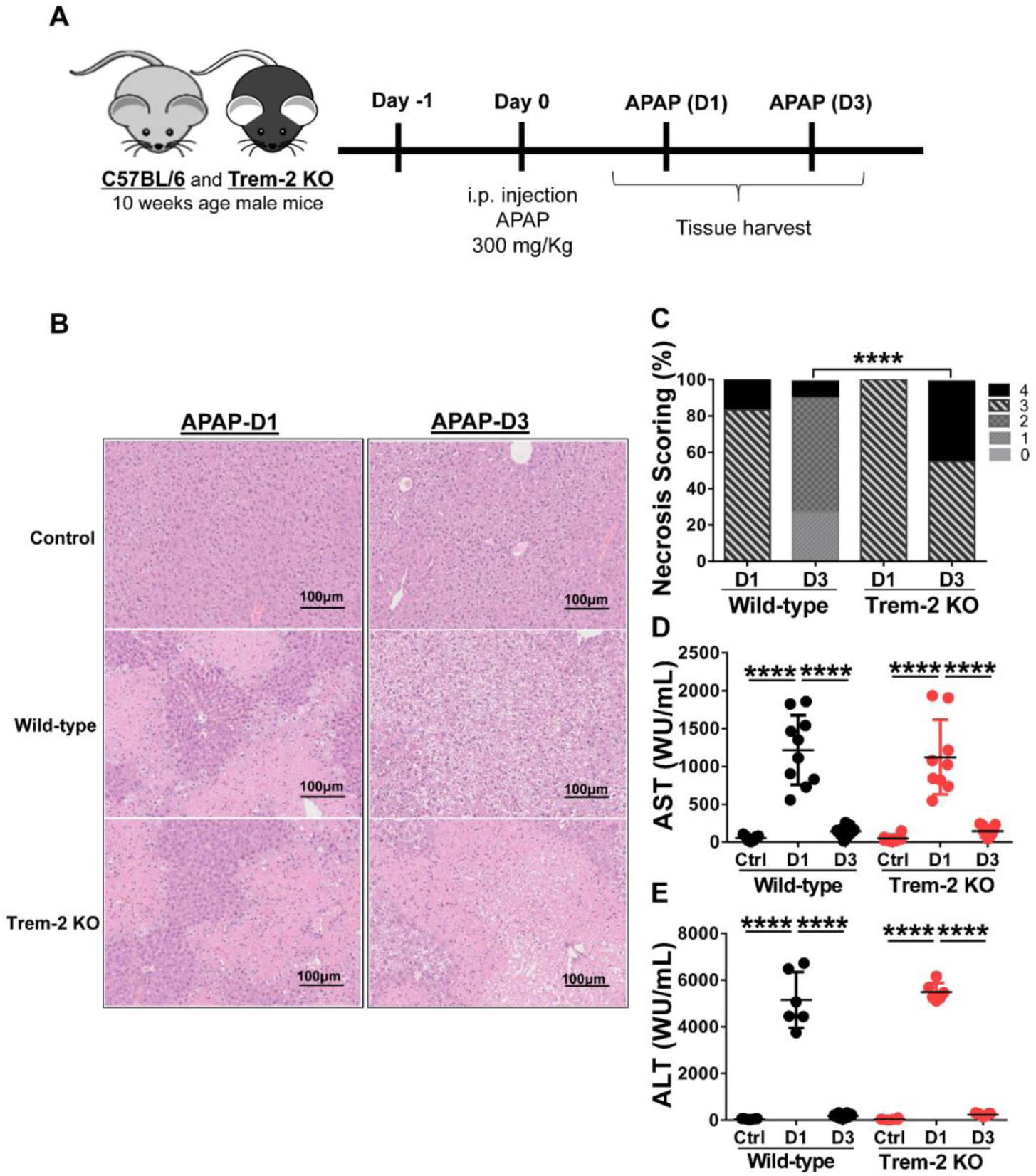
Trem-2 KO mice show impaired recovery from liver damage induced by APAP. Acute liver damage induced by single intra-peritoneal administration of acetaminophen (APAP) was assessed 1 day (APAP-D1) or 3 days (APAP-D3) post-injection of wild-type and Trem-2 KO mice (A). Hepatocyte necrosis was evaluated using hematoxylin-eosin staining (B) and scored from 0 to 4 according to location and extension of the necrotic lesions (C). Liver damage was assessed by quantification of hepatic enzymes AST (D) and ALT (E) in the serum. Mean values and standard deviations are represented. Statistics: Chi-square test in necrosis scoring (C); One-way ANOVA in (D); (n=6-11 mice/group). ****, p<0.0001.

### Trem-2 impacts on non-parenchymal cells dynamics during recovery from acute liver damage

Given the role of Trem-2 in macrophage functional activation^19^ we isolated non-parenchymal cells (NPCs) from APAP-treated mice at the time points of liver injury (D1) and tissue repair (D3) and performed a detailed analysis of the macrophage lineage cells populations **(Fig. 2A)**. Recruited hepatic macrophages (RHM) (CD45^+^ Ly6c^+^ F4/80^int^Mac1^high^) known to promote tissue inflammation^7,28^ were predominant at D1 but declined during the tissue repair phase (D3). RHM were found in similar proportions in wild-type and Trem-2 KO mice, suggesting that macrophage recruitment to the liver was not affected in absence of Trem-2 **(Fig. 2A and B)**. As expected, Kupffer cells (KCs) (CD45^+^ Ly6c^-^ F4/80^hi^ Mac1^int^) were highly represented in untreated mice and were severely reduced during injury (D1) in wild-type and Trem-2 KO mice **(Fig. 2A and B)**. However, we noted that in the tissue repair phase (D3) the recovery of KCs was slower in Trem-2 KO mice. We have recently shown that levels of CD26 enzymatic activity is a serum biomarker that mirrors severe reductions in KCs population^26^. Quantification of CD26 activity in the serum showed that at D3 Trem-2 KO mice reach slightly higher levels of CD26 activity **(Supplementary Fig. 1A)** corroborating the delayed KCs replenishment in Trem-2 KO mice.

**Figure 2.**
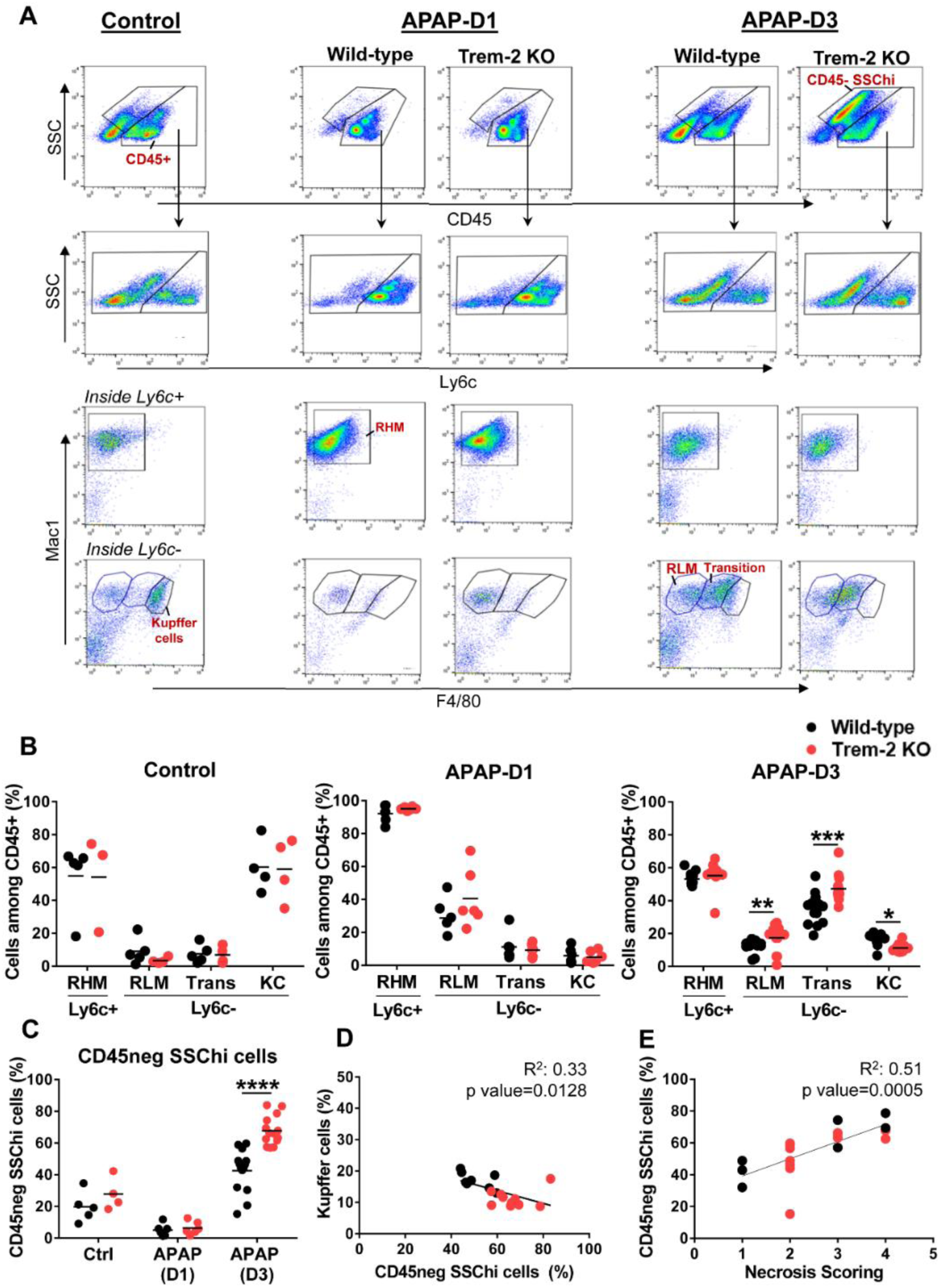
Impaired KC replenishment upon acute liver injury in Trem-2 KO mice and accumulation of non-hematopoietic CD45^neg^ SSC^hi^ cells. Flow cytometry analysis of liver non-parenchymal cells (NPCs) from wild-type and Trem-2 KO mice (control versus D1 versus D3) revealed distinct macrophage populations and a non-hematopoietic cell population (CD45^neg^ SSC^hi^). Among macrophages we identified Ly6c^+^ recruited hepatic macrophages (RHM, CD45^+^ Ly6c^+^ F4/80^low^ Mac1^+^) and Ly6c^-^ macrophages, namely, Kupffer cells (KC, CD45^+^ Ly6c^-^ F4/80^high^ Mac^int^); transition macrophages (Trans, CD45^+^ Ly6c^-^ F4/80^int^ Mac1^high^); and recruited-like macrophages (RLM, CD45^+^ Ly6c^-^ F4/80^low^ Mac1^+^) (A). Cell frequencies of macrophage populations (B) and non-hematopoietic CD45^neg^ SSC^hi^ population (C) in control, APAP-D1 and APAP-D3 wild-type and Trem-2 KO mice. Inverse correlation of KC frequency with CD45^neg^ SSC^hi^ population at APAP-D3 (D). Correlation of CD45^neg^ SSC^hi^ cells frequency with necrosis scoring at APAP-D3 (E). Symbols represent values from individual mice (n=6-11 mice/group). Group mean values are presented. Statistics: Two-way ANOVA in B and C. Pearson’s correlation test in D and E. *, p<0.05 **, p<0.01 ***, p<0.001 ****, p<0.0001.

Interestingly, our analysis revealed two related liver Ly6c^-^ macrophage populations with distinctive surface phenotypes: a CD45^+^ Ly6c^-^ F4/80^low^ Mac1^+^ population enriched at D1 and resembling the typical recruited Ly6c^+^ counterpart, herein named as recruited-like macrophages (RLM) **(Fig. 2A)** and a CD45^+^ Ly6c^-^ F4/80^+^ Mac1^hi^ population showing a surface phenotype close to KCs and named as transition macrophages that was strikingly enriched during tissue repair (D3) **(Fig. 2A and B)**. In contrast to KCs these populations accumulated in Trem-2 KO mice at D3 **(Fig. 2B)** indicating that in absence of Trem-2 the dynamics of macrophage hepatic repopulation was altered during the tissue repair response.

Using GFP-labeled monocyte transfers we show that after acute liver injury recruited bone-marrow monocytes give rise to RLM, transition macrophages and KCs **(Supplementary Fig. 2)**. Together these results suggest that in response to acute liver damage Trem-2 is a determinant of macrophage population dynamics promoting the replenishment of the KCs niche from recruited monocytes.

In addition, flow cytometry analysis uncovered a previously unnoticed non-hematopoietic CD45^neg^ SSC^hi^ population that emerges at D3 and accumulates in Trem-2 KO mice **(Fig. 2A and C)** paralleling the accumulation of transition macrophages. Strikingly, we noted that accumulation of non-hematopoietic CD45^neg^ SSC^hi^ cells inversely correlate with the KCs proportion in the livers of wild-type and Trem-2 KO mice **(Fig. 2D)** and directly correlated with the persistence of liver necrosis **(Fig. 2E)**. These findings suggest that in absence of Trem-2 imbalanced replenishment of liver macrophages associated with accumulation of transition macrophages results in overrepresentation of a non-parenchymal cell population, which correlates with impaired resolution of liver necrosis.

### Trem-2 ablation delays tissue repair and alters non-parenchymal cell dynamics in chronic liver damage

To extend our observations to chronic liver damage wild-type and Trem-2 KO mice were exposed to carbon tetrachloride (CCl4) treatment for 4 weeks and analyzed on day 1 (D1) and day 3 (D3) after treatment corresponding to established liver fibrosis and fibrosis regression time points, respectively **(Fig. 3A)**. At D1 Trem-2 KO mice showed a stronger fibrotic phenotype, with increased hepatocyte necrosis and fiber deposition **(Fig. 3B, C and D)**. Strikingly, fibrosis resolution response was compromised in Trem-2 KO mice by D3 as indicated by persistence of necrosis and collagen deposition that contrasted with nearly complete fibrosis regression in wild-type mice **(Fig. 3B, C and D)**. Accordingly, analysis of macrophage dynamics revealed that recruitment of macrophages at D1 was not affected in Trem-2 KO mice but at D3 KCs replenishment was impaired and non-hematopoietic CD45^neg^ SSC^hi^ cells were notoriously overrepresented **(Fig. 3E and F)**. Additionally, serum levels of CD26 activity tend to be higher at D3 in Trem-2 KO mice **(Supplementary Fig. 1B)**, strengthening the notion that KCs recovery was delayed in these mice. Similar to the acute model, CD45^neg^ SSC^hi^ cells accumulation at D3 was inversely correlated with KC proportions **(Fig. 3G)** and associated with the persistence of liver necrosis **(Fig. 3H)**. Furthermore, genetic expression of TNF-α in non-parenchymal cells (NPCs) correlated with the proportions of CD45^neg^ SSC^hi^ cell population **(Supplementary Fig. 3)**. These data reinforce that ablation of Trem-2 impacts the dynamics of macrophage populations favoring a pro-inflammatory milieu and accumulation of CD45^neg^ SSC^hi^ cells that are in turn associated with delayed resolution of tissue damage.

**Figure 3.**
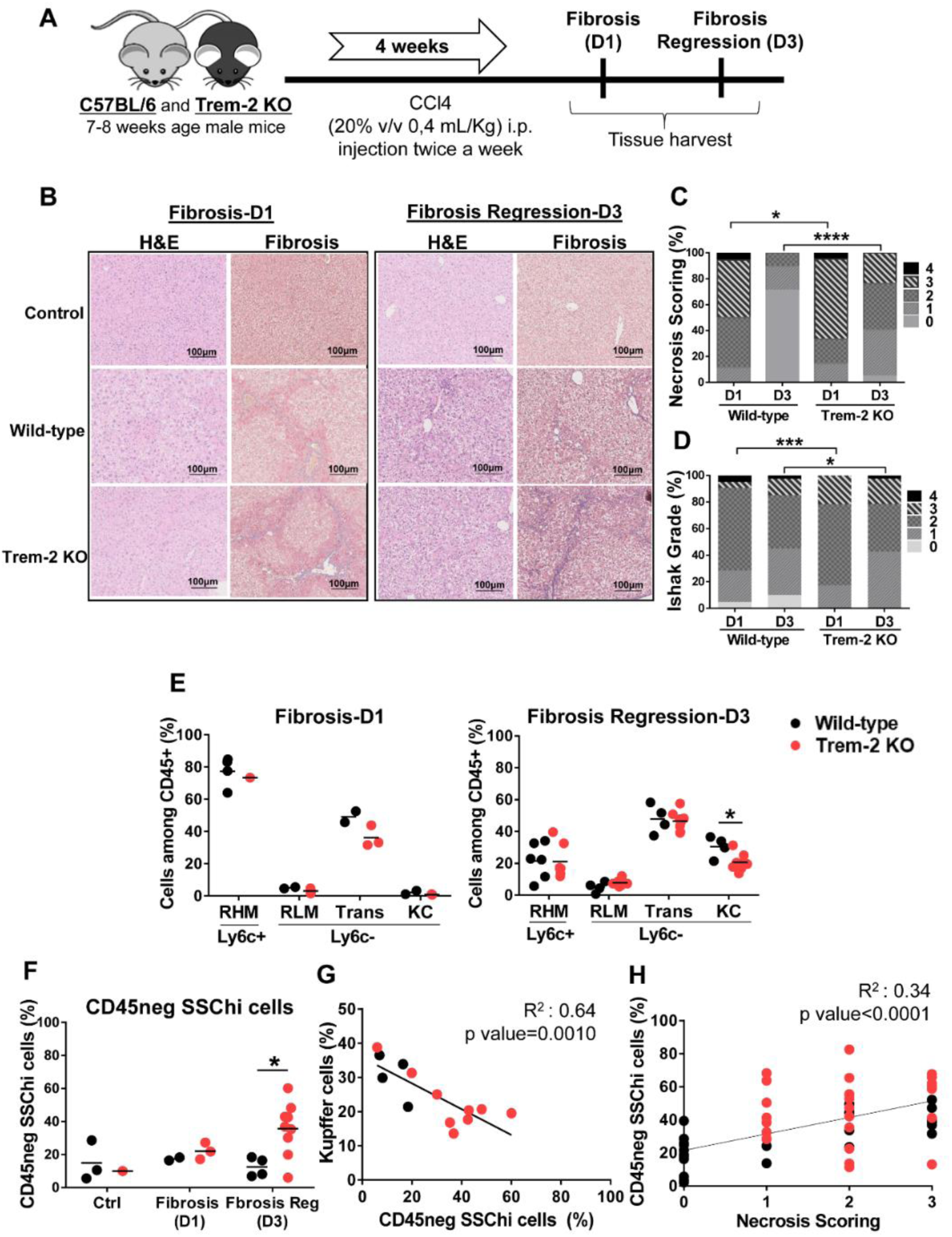
Responses to chronic liver injury are impaired in Trem-2 KO mice. Chronic liver injury was induced by administration of CCl4 during 4 weeks, twice a week. Mice were analyzed one or three days after the last injection, corresponding to fibrosis-D1 and fibrosis regression-D3 time points in wild-type mice (A). Liver necrosis was evaluated using hematoxylin-eosin (H&E) staining (B) and scored from 0 to 4 according to location and extension of the necrotic lesions (C). Masson’s trichrome (MT) staining was used to identify and quantify collagen fibers (blue) (B) using an adapted Ishak grading (D). Applying the flow cytometry criteria depicted in Figure 2 we quantified cell frequencies of distinct macrophage populations: recruited hepatic macrophages (RHM); Kupffer cells (KC); transition macrophages (Trans); recruited-like macrophages (RLM) (E) and of non-hematopoietic CD45^neg^ SSC^hi^ population (F) in wild-type and Trem-2 KO mice for control, fibrosis-D1 and fibrosis regression-D3 time points. Inverse correlation of KC frequency with CD45^neg^ SSC^hi^ population at Fibrosis Regression-D3 (G). Correlation of CD45^neg^ SSC^hi^ cells frequency with necrosis scoring at fibrosis regression-D3 time point (H). Symbols represent values from individual mice. Group mean values are presented. Statistics: For necrosis and Ishak grade scorings, groups were compared using Chi-square test (n=18-48 mice/group) (C). Two-way ANOVA in E and F. Pearson’s correlation test in G and H. *, p<0.05 ***, p<0.001, ****, p<0.0001.

### Trem-2 is upregulated in transition macrophages promoting acquisition of resident-like phenotype

Trem-2 RNA expression was quantified in sort-purified macrophage populations and non-hematopoietic CD45^neg^ SSC^hi^ cells from wild-type mice 3 days after APAP treatment (D3) **(Supplementary Fig. 4A)**. Remarkably, Trem-2 expression was upregulated in transition macrophages and almost undetectable in the other macrophage populations **(Fig. 4A)**. This strongly suggests that expression of Trem-2 in the transition macrophage population promotes adequate dynamics of KCs replenishment. Furthermore, Trem-2 was not expressed in CD45^neg^ SSC^hi^ cells **(Fig. 4A)** suggesting that their accumulation in Trem-2 KO mice may result from abnormal macrophage responses.

**Figure 4.**
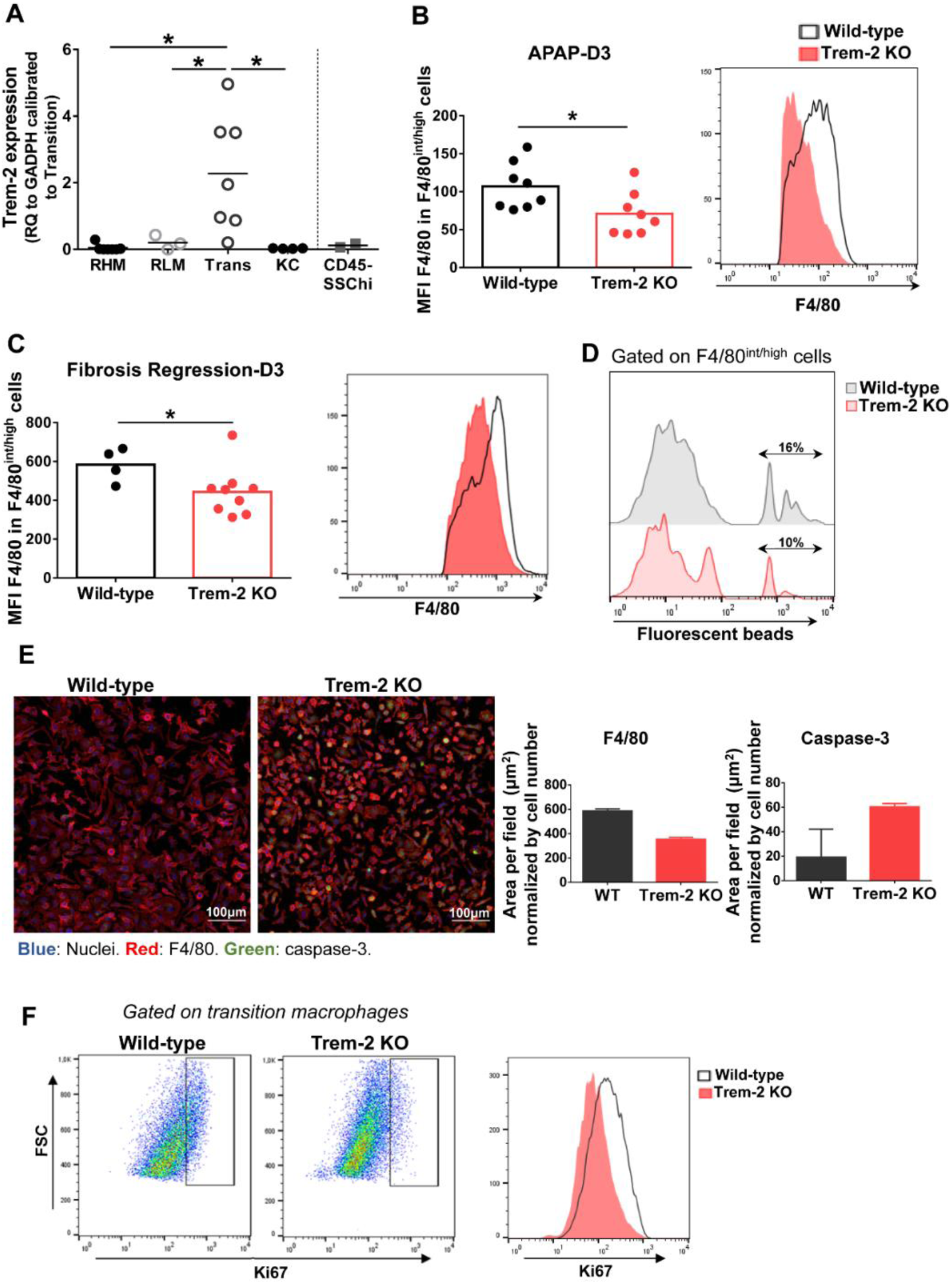
Trem-2 is implicated in acquisition of liver transition macrophages phenotype. Trem-2 gene expression was evaluated by qPCR in sort-purified macrophage populations and in non-hematopoietic cell population, CD45^neg^ SSC^hi^ in wild-type mice and represented as fold change to transition macrophages (A). F4/80 mean fluorescence intensity (MFI) among F4/80^int/high^ non-parenchymal cells from wild-type and Trem-2 KO mice at APAP-D3 (B) and fibrosis regression-D3 (C). Percent of F4/80^int/high^ non-parenchymal cells containing fluorescent beads in wild-type and Trem-2 KO at APAP-D3 one hour after i.v. injection of 50×10^6^ 2μm Yellow-Green fluorescent microspheres (D). Immunofluorescence of sort-purified APAP-D3 transition macrophages from wild-type and Trem-2 KO mice after 3 days of culture (Nuclei (blue), F4/80 (red), caspase-3 (green). Merged images are shown and quantification of F4/80 and caspase-3 stained area normalized by nuclei is plotted (E). Representative flow cytometry plots showing Ki67 staining in APAP-D3 transition macrophages from wild-type and Trem-2 KO mice (F). Statistics: One-way ANOVA and Tukey’s correction in A. Mann Whitney test in B and C. *, p<0.05.

Analysis of F4/80 surface expression, highly expressed in KCs, revealed that Ly6c^-^ macrophages from Trem-2 KO mice present decreased surface expression of F4/80 molecule upon acute **(Fig. 4B)** and chronic **(Fig. 4C)** injury. An *in vivo* phagocytosis functional assay with fluorescent beads at D3 after APAP treatment show that Trem-2 KO macrophages have decreased ability to phagocytose compared to wild-type **(Fig. 4D)**. Furthermore, we found that Trem-2 KO transition macrophages in culture maintain a rounder shape as assessed by the total F4/80 area **(Fig. 4E)**, while wild-type macrophages acquire a typical KC-like morphology. This suggests that Trem-2 is involved in the phenotype switch from transition to resident-like macrophage. Quantification of caspase-3 expression in culture and ki67 *ex vivo* staining in Trem-2 KO transition macrophages showed increased apoptosis and decreased proliferation **(Fig. 4E and F)**. These results indicate that ablation of Trem-2 impairs transition macrophage responses possibly by deterring phenotypic programming.

### Trem-2 drives transcriptional reprogramming in transition macrophages

To better discern the functional role of Trem-2 in macrophage phenotypic shifts we performed transcriptomic analysis in sort-purified macrophage populations of wild-type and Trem-2 KO mice **(Supplementary Fig. 4A)** at D3 after APAP treatment. Hierarchical algorithms and principal component analysis (PCA) clustered the different macrophage populations and clearly showed that the transcriptional profiles of RHM are in the vicinity of RLM and that transitional macrophages are closer to KCs **(Fig. 5A and Supplementary Fig. 5A)**. This is in agreement with accepted notions that recruited macrophages (RHM) home the liver as inflammatory macrophages and loose Ly6c expression subsequently giving rise to transition macrophages and KCs^29,30^.

**Figure 5.**
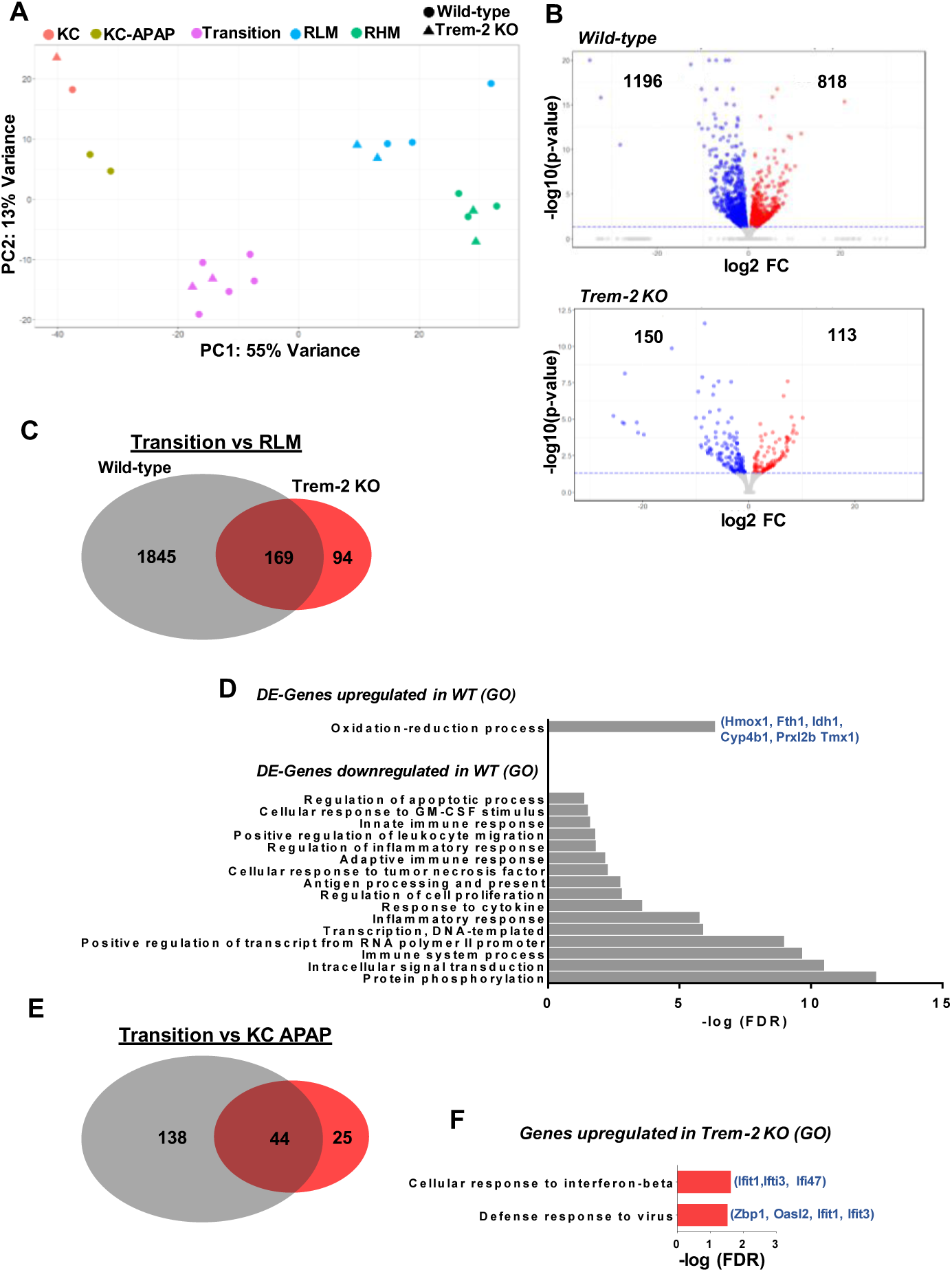
Transcriptomic profiling identifies a Trem-2 dependent program in transition macrophages. Principal component analysis (PCA) of transcriptomic data representing clustering of different sort-purified macrophage populations (represented by colors) from wild-type (circles) and Trem-2 KO mice (triangles) (A). Volcano plots representing differential expressed (DE) genes (q<0.05) between transition and recruited-like macrophages (RLM) in wild-type and Trem-2 KO mice. Red dots represent upregulated genes (LogFC>0), while blue dots represent downregulated genes (LogFC<0) (B). Venn diagram representing DE-genes between transition and RLM, which are common to wild-type and Trem-2 KO mice (middle), exclusively detected in wild-type (left) or in Trem-2 KO (right) (C). Gene Ontology (GO) enrichment analysis in the ‘Biological Process’ category for DE-genes upregulated or downregulated in transition versus RLM in wild-type mice (D). Venn diagram representing DE-genes upregulated in transition macrophages versus Kupffer cells (KC APAP-D3) in wild-type and Trem-2 KO mice (E). Gene Ontology (GO) enrichment analysis in the ‘Biological Process’ category for DE-genes exclusively upregulated in transition macrophages of Trem-2 KO mice (F). GO terms over-represented for FDR (Bejamini)<0.05.

On the other hand, wild-type and Trem-2 samples were clustered within each macrophage population **(Fig. 5A)**, suggesting that Trem-2 does not have a major impact in the global macrophage transcriptional programs. Given that Trem-2 affects the dynamical switching of transitional macrophage populations we performed a detailed analysis of differentially expressed (DE) genes comparing RLM and transition macrophages. We found that the transcriptional shift was more prominent in wild-type mice than in Trem-2 KO mice **(Fig. 5B and C)**. Interestingly, this shift encompassed the upregulation of genes associated to oxidation-reduction processes and downregulation of genes associated with inflammatory responses **(Fig. 5D)**, a pattern that was not observed in Trem-2 KO cells. We measured by qPCR the expression of two of these upregulated genes (Hmox1 and Fth1) in sorted transition macrophages confirming that activation of oxidative stress response mechanisms are blunted in Trem-2 KO transition macrophages **(Supplementary Fig. 6A)**. This analysis suggests that the transition macrophage transcriptional program was not fully acquired in Trem-2 KO cells.

Furthermore, comparison of transition macrophages and replenished KCs again showed that the transcriptional program switch is considerably less prominent in Trem-2 KO mice **(Fig. 5E)**. Gene ontology analysis revealed that genes specifically upregulated in transition macrophages from Trem-2 KO mice are associated to interferon-beta response, suggesting that these cells sustained a pro-inflammatory profile **(Fig. 5F)**. In addition, we analyzed the transcriptional switch between recruited macrophage populations comparing RHM to RLM and found a similar change extent in wild-type and Trem-2 KO mice **(Supplementary Fig. 6B)**. Moreover, gene ontology analysis showed that irrespective of Trem-2 expression, RLM have a transcriptional profile of increased proliferative capacity **(Supplementary Fig. 6C)**. Taken together these results show that Trem-2 expression in transition macrophages is key to shutdown the pro-inflammatory transcriptional program and increase resilience to oxidative stress during acquisition of resident macrophage functions.

### CD45^neg^ SSC^hi^ cells present an endothelial lineage transcriptomic profile

We next sought to discern the transcriptional program of the non-hematopoietic CD45^neg^ SSC^hi^ cells that accumulate in the liver in response to tissue injury and correlate with reduced liver KCs and persistent liver damage (**Fig. 2D-E and 3G-H**). The transcriptomic profiles of sorted CD45^neg^ SSC^hi^ cells in APAP-treated (D3) and untreated mice **(Supplementary Fig. 4B)** were closely related when using KCs as a reference population **(Fig. 6A and Supplementary Fig. 5B)**. Gene ontology analysis performed for the DE-genes common to control and APAP-D3 CD45^neg^ SSC^hi^ cells showed striking enrichment in terms related to endothelial cell identity and function **(Fig. 6B)**. In addition, DE-genes upregulated in CD45^neg^ SSC^hi^ cells of APAP–D3 treated mice revealed enrichment in pathways related to epithelial to mesenchymal transition and to regulation of Wnt signaling pathway **(Fig. 6C)**. On the other hand, DE-genes up-regulated in CD45^neg^ SSC^hi^ cells from control mice revealed enrichment in functional pathways involved in blood coagulation, hemostasis and fibrinolysis, typical of endothelial cells **(Fig. 6D)**. These transcriptomic data clearly identified the CD45^neg^ SSC^hi^ population accumulating in the damaged liver as belonging to the endothelial cell lineage leading us to operationally name these cells as Liver Damage-associated Endothelial Cells (LDECs). In addition, the ontology analysis suggests that LDECs are undergoing endothelial de-differentiation.

**Figure 6.**
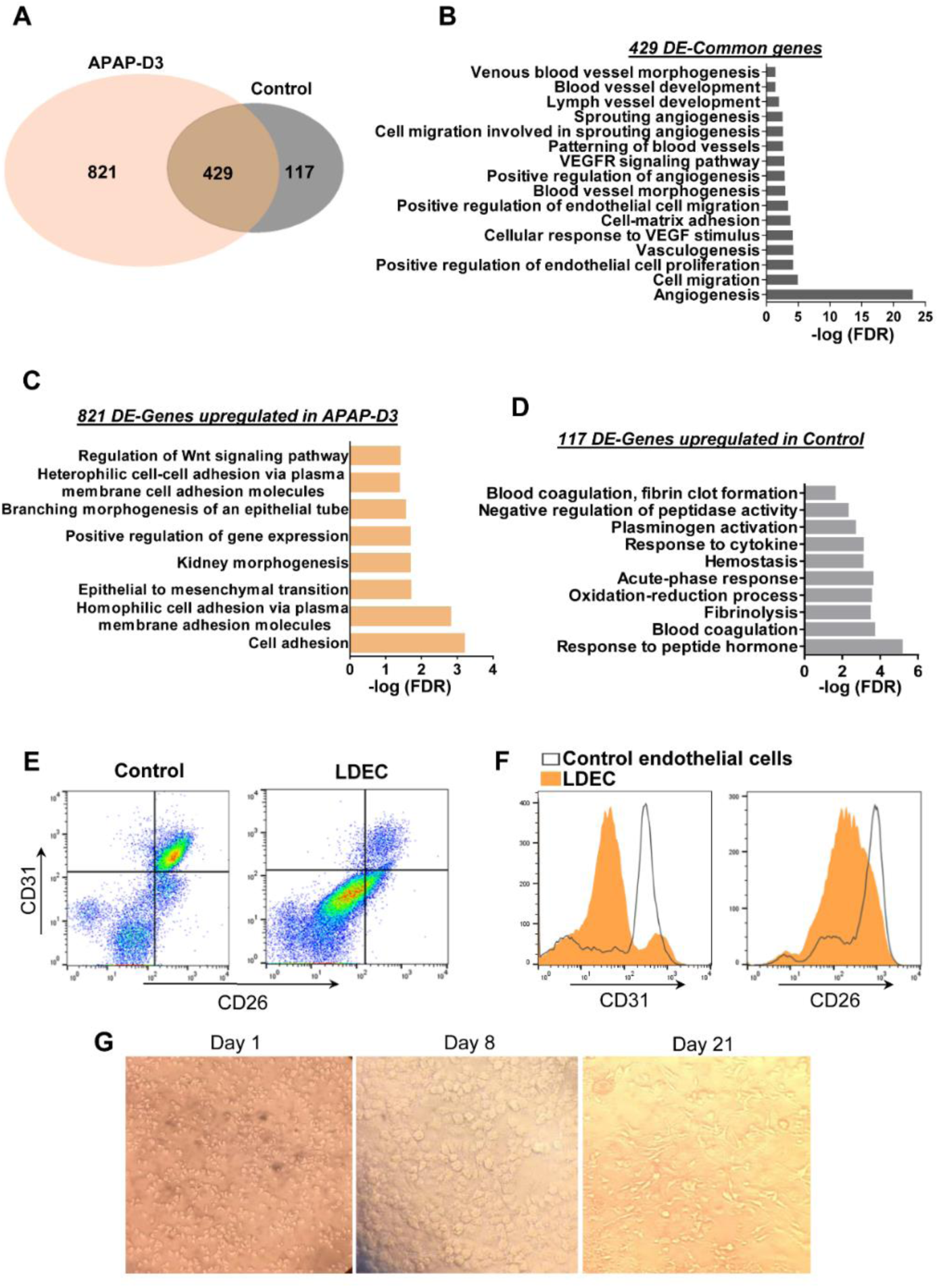
Liver damage-associated endothelial cells identified among the CD45^neg^ SSC^hi^ population by transcriptomic and phenotypic analysis. Venn diagram representing differentially expressed DE-genes upregulated in CD45^neg^ SSC^hi^ population from APAP-D3 and untreated control mice using Kupffer cells (KC APAP-D3) as a reference (A). Gene Ontology (GO) enrichment analysis in the ‘Biological Process’ category for common DE upregulated genes in APAP-D3 and control compared to KCs (B) for DE upregulated genes exclusively in APAP-D3 (C) and genes upregulated exclusively control mice (D). GO terms over-represented for FDR (Bejamini)<0.05. Flow cytometry plots showing double staining for CD31 and CD26 in APAP-D3 (LDECs) and control CD45^neg^ SSC^hi^ cells (E, F). Sort-purified LDECs cultured in M-CSF supplemented medium at 1, 8 and 21 days (G).

### LDECs proliferate and differentiate in vitro

We characterized LDECs by flow cytometry using liver endothelial cell surface makers, namely CD31 and CD26. Strikingly, LDECs express lower levels of these markers as compared to endothelial cells from control mice **(Fig. 6E and F)**. This is line with previous reports showing that endothelial cells undergoing endothelial to mesenchymal transition downregulate endothelial specific markers such as CD31^31^. We next explored the ability of sort-purified LDECs to differentiate *in vitro* in presence of macrophage-colony stimulating factor (M-CSF), a growth factor able to promote angiogenesis^32^. LEDCs were very small and granulous until day 8 when larger sized, round-shaped cells emerged and expanded in the culture, eventually developing typical endothelial morphology by 21 days **(Fig. 6G)**. These results reveal that LDECs are at a particular activation state characterized by down-modulation of endothelial cell markers and ability to differentiate into morphologically distinct cells, indicative of their precursor potential.

## Discussion

This work revealed that Trem-2 controls the replenishment of liver macrophage populations after acute and chronic hepatotoxic damage and is a critical determinant of swift tissue repair responses conditioning the emergence of endothelial lineage cells during regeneration.

A recent report proposed that lack of Trem-2 expression in non-parenchymal cells contributes to increased fibrosis in Trem-2 KO mice submitted to CCl4 treatment and to increased liver inflammation after acute APAP treatment^24^. However, we noted that Trem-2 KO mice did not show significantly higher intensity of liver damage during acute injury (APAP-D1) but showed sustained tissue damage throughout the tissue repair phase (APAP-D3). Likewise, Trem-2 KO mice presented slightly heightened necrosis during the inflammatory phase of chronic liver injury (CCl4-D1) and a marked delay in subsequent resolution of tissue necrosis and fibrosis (CCl4-D3). While Trem-2 may influence the initial hepatic inflammatory and fibrotic reactions^24^, here we focused on its impact in liver macrophages responses during recovery from drug-induced damage.

Macrophage phenotypic plasticity is well illustrated in the responses to liver tissue damage^8,9^. In particular, we identified a transition macrophage population that expresses Trem-2 in high levels and predominates during the recovery phase from acute and chronic damage. These cells are derived from circulating monocytes and both transcriptional and phenotypic profiling placed them as the immediate source of resident-like macrophages in the recovered liver. These observations strongly suggest that Trem-2 signaling plays a role in the dynamics and efficiency of this phenotypic transition explaining the delayed replenishment of the KCs compartment in Trem-2 KO mice.

Noteworthy, a Trem-2 dependent transcriptional program has been associated with emergence of restorative macrophage populations in the context of brain tissue degeneration and adipose tissue inflammation^21,22^. Similarly, liver transition macrophages show upregulation of genes associated with this Trem-2 dependent transcriptional signature **(Supplementary Fig. 7)**. Thus, our results contribute to identify Trem-2 as a critical determinant of macrophage plasticity required for a swift recovery from tissue damage.

Acquisition of the transition phenotype in wild-type macrophages includes down-modulation of pro-inflammatory genes and up-regulation of genes involved in oxidative stress responses, not observed in Trem-2 KO macrophages. Redox regulation is critical to cellular stress control mechanisms^33^. Accordingly, transition macrophages from Trem-2 KO mice show decreased survival and proliferative capacities. These observations are in line with reports of reduced survival of Trem-2 KO microglia cells during neurodegenerative processes^34,35^. Furthermore, reactive oxygen species (ROS) were increased in Trem-2 KO bone-marrow derived macrophages and hepatic lipid peroxides were increased during liver damage in Trem-2 KO mice^24^. Together, these findings suggest that transition macrophages may play a relevant role in ROS clearance in the damaged liver.

Remarkably, ablation of Trem-2 leads to increased accumulation of a non-hematopoietic population that we identified as an endothelial cell population herein named as Liver Damage Endothelial Cells (LDECs). This population appeared during tissue repair phases and its accumulation correlated with the severity of tissue damage. It has been recently proposed that vascular endothelial stem cells residing in the liver are activated upon acute liver injury and act as angiogenesis-initiating cells showing remarkable vascular regenerative capacity^36^. On the other hand, a specific subset of liver sinusoidal endothelial cells was found to sustain liver regeneration after hepatectomy by releasing angiocrine trophogens^37^. Nevertheless, the development and rules of engagement of endothelial progenitors in liver angiogenic repair responses remain unclear.

Interestingly, LDECs transcriptional profile denotes a differential functional activation status as compared to control endothelial cells. Gene ontology analysis revealed that LDECs are involved in biological processes related to epithelial to mesenchymal transition (Fig. 2.6C). A similar process designated endothelial do mesenchymal transition (EndMT) has been reported to play important roles in pathogenesis of many diseases^38^ as well as in regenerative processes^31^. EndMT promotes cell de-differentiation giving rise to mesenchymal stem cells with the ability to differentiate into new cell types^39^. These cells were shown to differentiate into endothelial cells contributing for neovascularization^38^.

We noted that accumulation and persistence of LDECs in Trem-2 KO mice is counterweighed by a slower replenishment of the KCs compartment suggesting that dynamics of macrophage repopulation may regulate LDECs appearance and/or function. Previous studies reported interactions between macrophages and endothelial cells undergoing EndMT^40,41^. In particular, in an experimental model of atherosclerosis it was shown that macrophages have the ability to partially induce this transition^40^. Interestingly, M-CSF, an essential regulator of macrophage development, was able to induce marked LDECs morphological changes *in vitro*, highlighting their de-differentiated state and intrinsic proliferative and differentiation potential.

In sum, this work describes Trem-2 as a promotor of macrophage phenotypic switching during tissue repair by shutting down the recruited macrophage inflammatory profile, enhancing oxidation-reduction responses and allowing KCs replenishment. In parallel, we identified an endothelial cell population (LDECs) that emerges during tissue repair with a distinct transcriptional profile and phenotypic features of endothelial de-differentiation. In conclusion, Trem-2 KO mice revealed that the dynamics of pro-restorative macrophage populations during repair responses is interlinked with emergence of LDECs and delayed recovery from tissue damage.

## Acknowledgements

The authors acknowledge the histology, flow cytometry, genomics, antibody production and bioinformatics units at IGC, in particular Dr. Rui Pedro Faísca for excellent technical assistance and Alexander Marta for analysis of immunofluorescence images.

## Notes

Financial Support This work was developed with the support of the research infrastructure Congento, project LISBOA-01-0145-FEDER-022170, co-financed by Lisboa Regional Operational Programme (Lisboa 2020), under the Portugal 2020 Partnership Agreement, through the European Regional Development Fund (ERDF), and FCT - “Fundação para a Ciência e a Tecnologia” (Portugal). This work was partially supported by ONEIDA project (LISBOA-01-0145-FEDER-016417) co-funded by FEEI - “Fundos Europeus Estruturais e de Investimento” from “Programa Operacional Regional Lisboa 2020” and by national funds from FCT through grants PTDC/BIM-MET/2115/2014, PTDC/BIM-MET/4265/2014 and iNOVA4Health (UID/Multi/04462/2013). IC was supported by a FCT fellowship PD/BD/105997/2014.

