## Supplementary materials and figures for "Trem-2 promotes emergence of restorative macrophages and endothelial cells during recovery from hepatic tissue damage"

**Supplementary Materials and Methods**

***CD26/DPP-4 enzymatic activity***

Serum from total blood was measured for DPP-4 activity using a fluorimetric assay with 200 uM of the substrate H-Gly-Pro-AMC HBr (#I-1225, Bachem, Bubendorf, Switzerland). Measurements at excitation/emission 360/460 nm were taken every 5 minutes during one hour and the slope of fluorescence was used as measure of cleaved substrate by CD26/DPP-4, reported as arbitrary units.

***Monocyte transfers***

To track recruited macrophages in the liver we isolated monocytes from the spleen of B6.Actin-GFP mice as previously described[1]. Briefly, spleens were collected, smashed and red blood cells lysed. Cells were firstly stained with biotinylated anti-TCR-β, anti-B220 and anti-NK1.1 antibodies and depleted with anti-biotin microbeads using MACS LD columns (MACS- Miltenyl Biotec). GFP-labeled monocytes, identified as $Mac1^+$ and $Ly6c^+$ by flow cytometry were counted and $2.5x10^6$ cells were injected intravenously in wild-type recipients. (See antibodies table below).

***Image Analysis***

Quantification of F4/80 fluorescence was obtained by thresholding each stitched tile using the Mean Auto-Threshold algorithm of FIJI software package[2] and measuring the total area of F4/80 normalized by the number of nuclei to control for cell confluency. Total number of nuclei was determined by thresholding the DAPI channel using the Otsu Auto-Threshold algorithm and counting them through FIJI particle analyzer functions.

Similar methods were employed to quantify Caspase-3 fluorescence, applying the Moments Auto-Threshold and dividing total area by number of nuclei for normalization. Total number of nuclei was obtained by making each stitched tile binary in the DAPI channel, applying the Watershed function, and recording total Count.

***RNA Sequencing analysis***

***Quality Assessment and Alignment.*** Prior to alignment, quality of the sequences was assessed using FASTQC[3] and MultiQC[4]. Alignment was performed against the *Mus musculus* genome version 95, with the annotation file for the genome version 95, both obtained through the Ensembl website and using STAR[5], with default parameters and with the option of *GeneCounts*.

***Data Analysis.*** The files obtained from *GeneCounts* option were imported to R, taking into account the strandness inherent to the sequencing protocol. Downstream analysis was performed using DESeq2 (version 1.22.2)[6]. Data from raw counts used to create the Principal Component Analysis plot and Heatmaps were normalized through a Variance Stabilizing Transformation (VST)[7]. The log2FC provided by the standard DESeq2 model was shrunk using the 'ashr' option[8]. Gene Information was obtained using the package *org.Mm.eg.db*. For the purposes of this study, genes were considered differentially expressed when the p-value, adjusted using false discovery rate (FDR), was below 0.05. Log2FC tables of differentially expressed genes (DE-genes) are available at https://figshare.com/s/3684f039abc8c96313a3. Gene ontology analysis was performed using DAVID Bioinformatics Resources 6.8 webtool (https://david.ncifcrf.gov/) and enriched terms considered for FDR<0.05 using Benjamini Hochberg method.

***Antibodies***

| **Immunohistochemistry/ Immunofluorescence** | | |
|---|---|---|
| **Antibody** | **Supplier** | **Clone** |
| Rat anti-F4/80 | BioRad | Cl:A3-1 |
| Rabbit anti-Caspase-3 | Abcam | polyclonal |
| Goat anti-rat IgG Alexa Fluor 647 | ThermoFisher | polyclonal |
| Goat anti-rabbit IgG Alexa Fluor 568 | ThermoFisher | polyclonal |
| **Flow Cytometry** | | |
| **Antibody** | **Supplier** | **Clone** |
| Anti-mouse-Fc-block/CD16/32 | IGC (antibody facility) | 2.4G2 |
| Anti-mouse PE CD45 | BioLegend | 104.2 |
| Anti-mouse Alexa-700 F4/80 | BioLegend | BM8 |
| Anti-mouse efluor450 Ly6c | BioLegend | HK1.4 |
| Anti-mouse Brilliant Violet-785-CD11b/Mac-1 | eBiosciences | M1/70 |
| Anti-mouse Alexa-488 Ki67 | BD Pharmigen | B56 |
| Anti-mouse Fitc CD26 | BioLegend | H194-112 |
| Anti-mouse APC CD31 | BD Pharmigen | MEC 13.3 |

| | | |
|---|---|---|
| Anti-mouse biotinylated TCR beta | IGC (antibody facility) | H57-597 |
| Anti-mouse biotnylated B220 | IGC (antibody facility) | RA3-6B2 |
| Anti-mouse biotinylated NK1.1 | IGC (antibody facility) | PK136 |

### Supplementary Figures

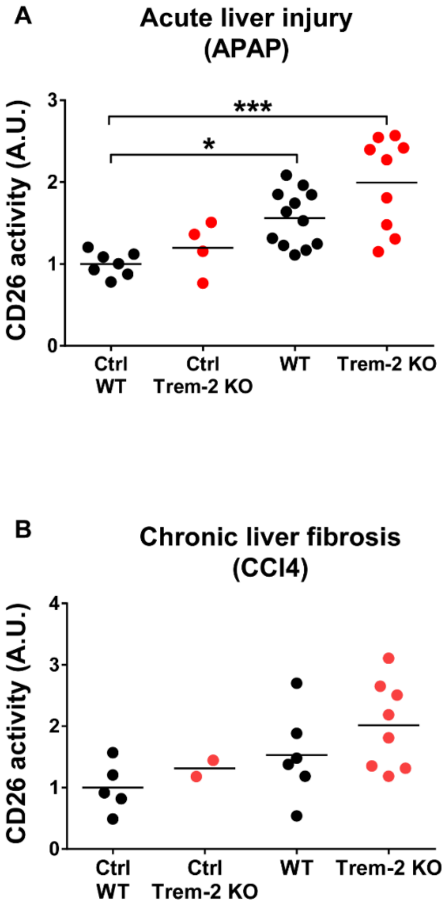


**Supplementary Figure 1. Serum CD26 activity during recovery from acute liver damage and fibrosis regression.** CD26 activity represented in arbitrary units (A.U.) was measured in the serum of wild-type and Trem-2 KO mice, during recovery from APAP (D3) (A) and during fibrosis regression (D3) induced by CCl4 (B) or in mice left untreated (Ctrl). Symbols represent individual mice. One-way ANOVA *, p<0.05 ***, p<0.001.

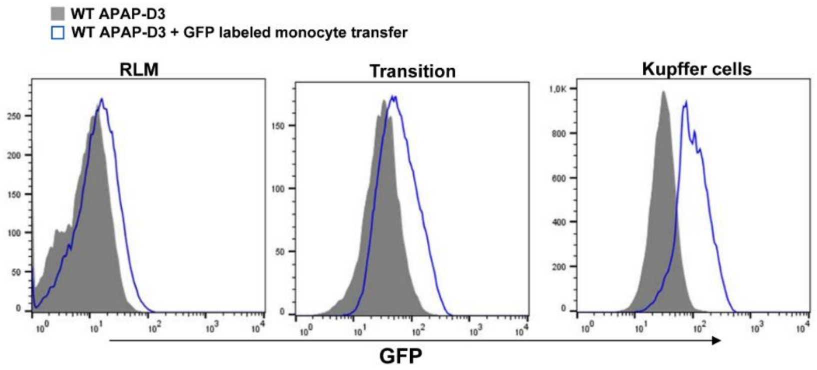


**Supplementary Figure 2. Recruited monocyte tracking in liver macrophage populations following acute liver damage.** Monocytes from B6.Actin-GFP mice were intravenously injected 12 hours after APAP treatment in wild-type mice and non-parenchymal cells were isolated at APAP-D3. Flow cytometry histograms show enrichment in $GFP^+$ cells compared to non-transferred mice in different macrophage populations: RLM, Transition and Kupffer cells.

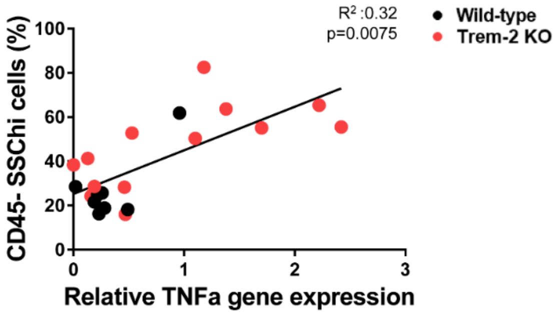


**Supplementary Figure 3. Accumulation of CD45$^{neg}$ SSC$^{hi}$ cells correlates with expression of pro-inflammatory cytokines in the liver.** Positive correlation between frequency of CD45$^{neg}$ SSC$^{hi}$ population and TNFα gene expression in liver non-parenchymal cells during recovery from chronic injury (CCl4-D3). Statistics: Pearson's correlation test.

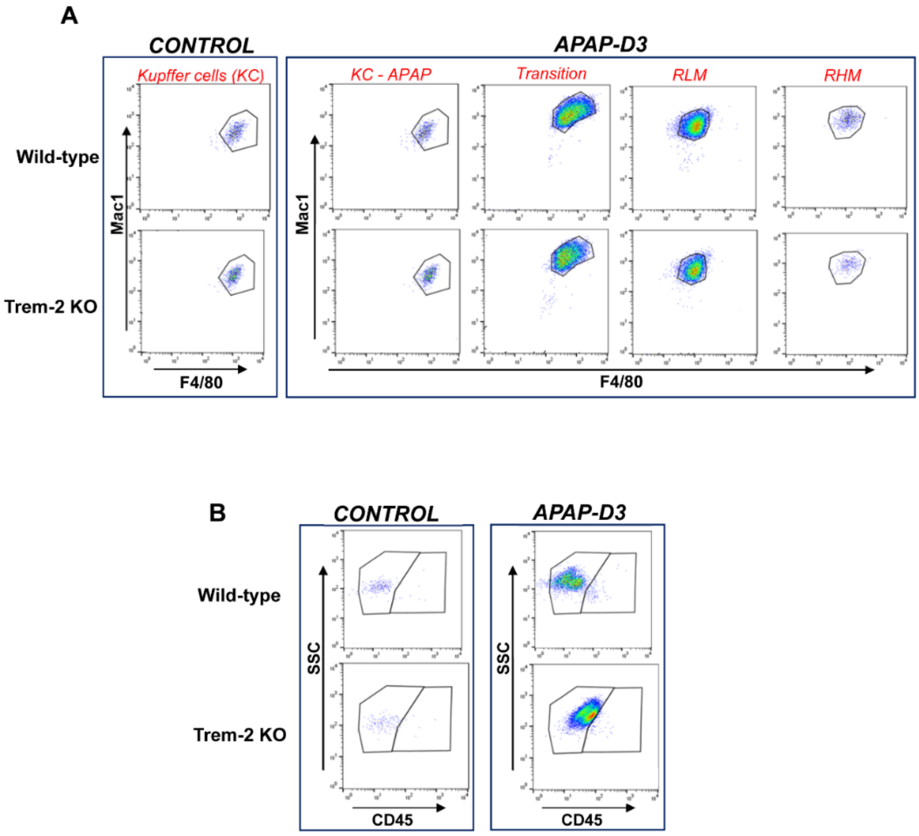


**Supplementary Figure 4. FACsort-purified macrophage populations and CD45$^{neg}$ SSC$^{hi}$ cells for transcriptomic analysis.** Wild-type and Trem-2 KO mice received a single intraperitoneal injection of APAP and macrophage populations and CD45$^{neg}$ SSC$^{hi}$ cells were sorted at APAP-D3 and in control mice. Macrophage populations were identified using CD45, Ly6c, Mac1 and F4/80 markers and the purity of each population after sorting is shown (A). CD45$^{neg}$ SSC$^{hi}$ population was identified using CD45 marker in APAP-D3 and in control mice and the purity after sorting is shown (B). Each sample for transcriptomic analysis was obtained from pools of 4 mice.

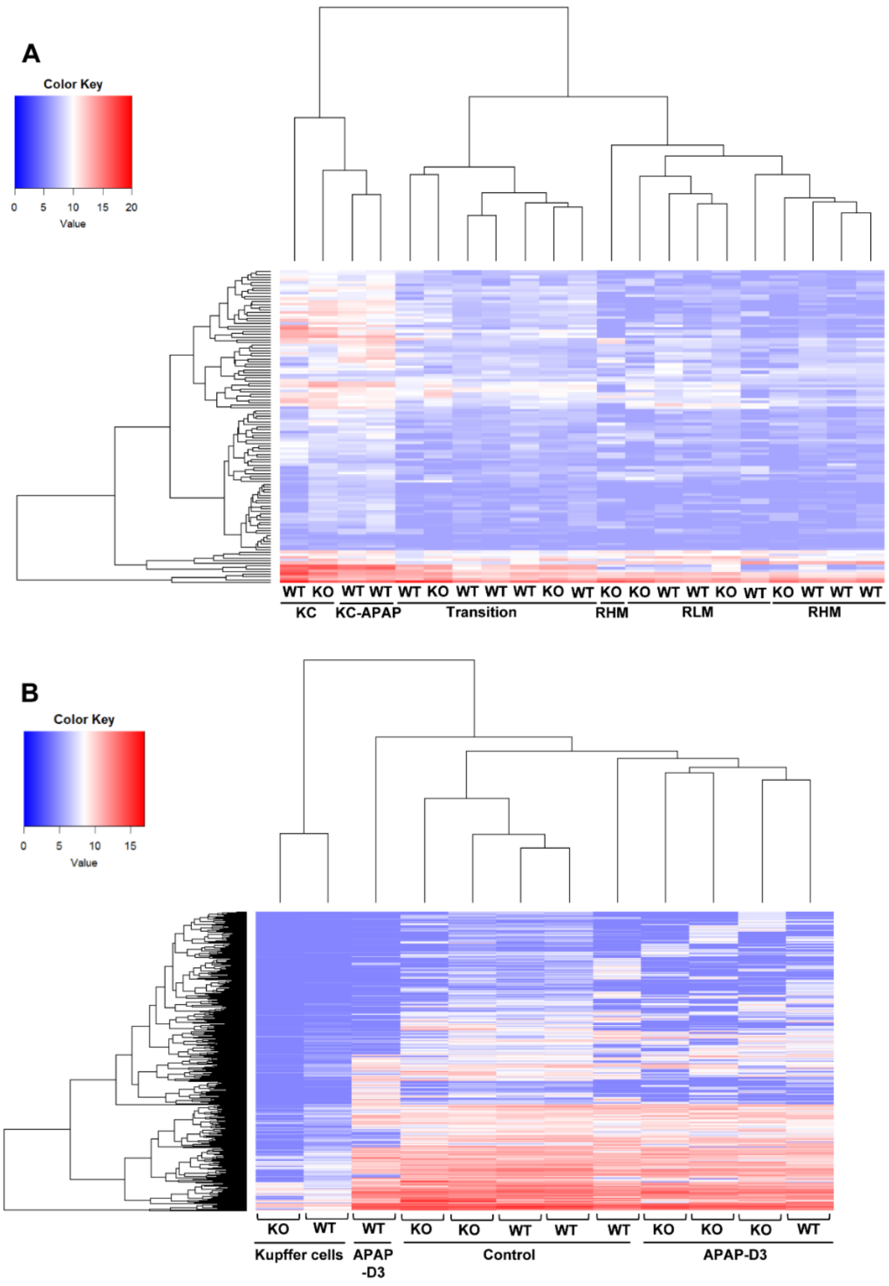


**Supplementary Figure 5. Heatmaps of DE-genes representing hierarchical clustering of macrophage and CD45$^{neg}$ SSC$^{hi}$ cell populations.** Heatmap represents 100 KCs associated genes[42] for different macrophage populations (A). Heatmap showing differentially expressed (DE) genes in CD45$^{neg}$ SSC$^{hi}$ cells using KCs as reference: 821 genes upregulated in APAP-D3, 117 upregulated in control and 428 common to APAP-D3 and control as identified in Fig. 2.6A (B).

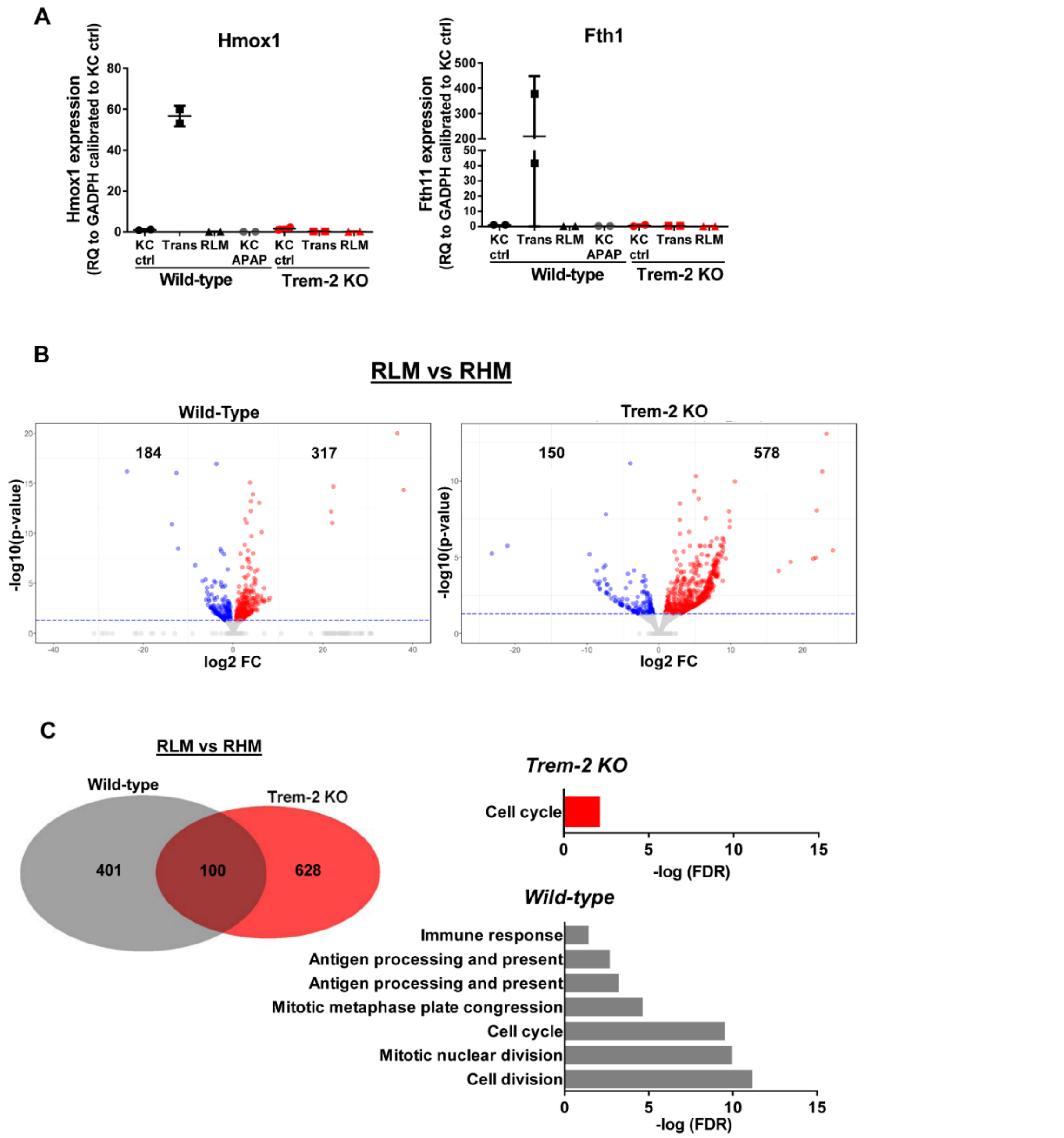


**Supplementary Figure 6. Gene expression and transcriptomic analysis of macrophage populations.** Hmox1 and Fth1 gene expression was evaluated by qPCR in sort-purified KCs, transition and RLM populations from wild-type and Trem-2 KO mice at APAP-D3 (A). Volcano plots representing differential expressed (DE) genes (q<0.05) between RLM and RHM in wild-type and Trem-2 KO mice. Red dots represent upregulated genes with LogFC>0, while blue dots represent downregulated genes with LogFC<0 significant for q<0.05 (B). Venn diagram representing DE-genes between RLM and RHM which are common to wild-type and Trem-2 KO (middle), exclusive for wild-type (left) or Trem-2 KO (right). Gene Ontology (GO) enrichment analysis in the 'Biological Process' category for DE-genes in wild-type mice and Trem-2 KO mice (C).

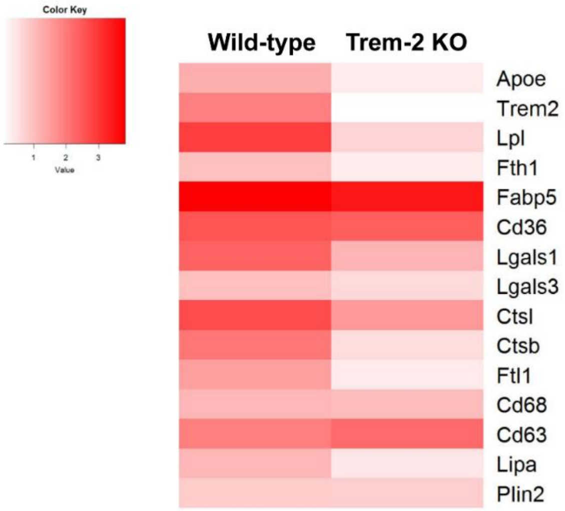


**Supplementary Figure 7. Heatmap illustrating genes regulated by Trem-2 in transition macrophages.** Log fold change (logFC) of genes previously associated to Trem-2 transcriptional signature[21,22]. Heatmap represents DE-genes upregulated in transition versus RLM at APAP-D3 in wild-type and Trem-2 KO mice.
